# Estimation of Genetic Diversity in Genus Mentha Collected From Azad Jammu And Kashmir, Pakistan

**DOI:** 10.1101/866111

**Authors:** Fozia Abasi, Israr Ahmad, Sami Ullah khan, Khawaja Shafique Ahmad, Aneela Ulfat, Rubab Khurshid

## Abstract

Mints are perennial aromatic herbs used both for medicinal and aromatic purposes. Flora of Pakistan has reported six species of genus *Mentha.* Taxonomy of genus *Mentha* is more complex and confusing due to inter specific hybridization. The present research is the first documented report from Pakistan for the purpose to dissect *Mentha* specimens using molecular tools. A total of 17 SCoT and SSR markers used to dissect genetic diversity among 41 *Mentha* specimens. The results revealed substantial variation among *Mentha* specimens. The molecular data analyzed through NTSYS and Power marker software’s. Dendrogram constructed on the base of similarity coefficients generated using weighted pair group method of arithmetic means (UPGMA) recorded high level of polymorphism. Polymorphic Information Content (PIC) for molecular markers recorded in the range of 5-8. Mean genetic distance (GD) was estimated in the range from 0.35% to 100%. The minimum and maximum GD recorded in one combination each as P2-P4 and M41-P10. The present study was explored the efficiency of SCoT and SSR markers for evaluating the genetic diversity of medicinal plants. The present research was concluded that both morphological and molecular dendrogram determined considerable level of diversity among *Mentha* species. Furthermore, specific primers and DNA chloroplast technology could be needed for further molecular analysis to refine the data more up to varietal level.

## INTRODUCTION

Genus Mentha belong to family Lamiaceae, comprises of twenty five to thirty species found worldwide [1]. The taxonomy of this genus is more complicated, more than three thousand names from species to formae distributed consequently to modern plants taxonomy [2]. The taxonomy of Mentha is ambiguous due to continuous polyploidy and interspecies hybridization that occurring in wild and cultured population [3]. Genus Mentha found in all five continents, extensive distribution and native to north temperate regions. On the basis of morphological variations genus Mentha reproduced on different taxonomic rank and documented to this genus during the past 200 years. Due to hybridization *Mentha* species showed complex variations among the wildest populations [4]. Different species of this genus is described on the basis of morphology of inflorescence Linnaeus. In addition to morphological traits, molecular markers and cytological tools has been used to assess the genetic variation, taxonomic and phylogenetic relationships among and within the *Mentha* species [1]. On the basis of morphogenetic tools, six common species of genus Mentha reported from Pakistan i.e. M. *Pulegium, Arvensis, M. spicata, M.longifolia, M.piperita* and *M. royleana* [5].

Mint produce important secondary metabolites that forage poisonous free radicals, check and explored potential sources for natural antioxidants due to the free radicals scavenging activity in *Mentha species* [6]. Due to presence of volatile compounds some typical secondary metabolites it shows cytotoxic and antioxidant activities [7] [8] [9]. It have a substantial significance in the botanical economy and to the pharmaceutical industry, mainly because of presence of essential oils and their antimicrobial properties, used since ancient times for the treatment of several digestive tract diseases and in culinary [10]. Genus *Mentha* species showed antioxidant, anti-inflammatory, anti-microbial activity, and anti-cancer activities. In *Mentha* species secondary metabolites terpenes mostly present in leaf epidermal glands, stem and propagative, indeed use for both medicinal and aromatic purposes and their flowers, leaves, stem are used as flavorings in commercial spice, flavor and herbal teas [11] [12]. Mint has pharmaceutical properties, therefore consumed for treatment of different diseases like bronchitis, flatulence, anorexia and consumption have astonishing due to bulk of uses, it also have great economical values and their essential oils are used like confection, cosmetics, oral sanitation products, medications, insecticides, as a fragrance enhancing mediator in toothpastes [13] [14].

Morphological tools i.e. leaf area, inflorescence color, and shape of leaf among the oldest markers used to identified Mentha species, are not informative to evaluate the genetic differences, because of genetic diversity in Mentha species due to environmental changes that cause variations in phenotype of individuals [15]. Understanding the genetic multiplicity of *Mentha* species is required for crops improvement [14]. Morphological traits, cellular biochemical, and isozyme polymorphism has been used to detect the genetic variability among the *Mentha* specimens [16]. The molecular markers are actual tools for identification of genotype similarity through DNA fingerprinting and used to check change at genetic level due to environmental effects [17] [2]]. AFLP marker were used for inter-specific diversity among Mentha hybrid [3].

Due to high polymorphism in morphology and great diversity in essential oil composition genus needs careful taxonomic reassessment with modern techniques. Therefore, the present research was design to estimate genetic diversity among *Mentha* species through molecular markers. In the present research, forty one Mentha specimens were studied by using molecular and morphological markers.

## MATERIAL AND METHODS

### Study Area

Azad Jammu & Kashmir state falls within the Himalayan orogenic belt. Topography of Azad Kashmir is mainly hilly and mountainous categorized by deep ravines, rugged, and undulating terrain. Northern districts (Neelum, Muzaffarabad, Hattian, Bagh, Haveli, Poonch, and Sudhnoti) are generally mountainous while the southern districts (Kotli, Mirpur, and Bhimber) are comparatively plain. The ecosystems mountain are relatively unstable and have low intrinsic productivity, within this fragile environment there is a great diversity of ecological niches upon which people base their livelihood. The area is full of natural beauty with thick forest, fast flowing rivers and winding streams and main rivers are Jhelum, Neelum and river Poonch [18].

### Plant material

A total of 41 *Mentha* specimens collected from different geographical zones of Azad Jammu & Kashmir as shown in Table 1. All specimens pressed, dried and paste on standard herbarium sheet. The specimens were identified with the help of flora of Pakistan and other available literature. The identified specimens were submitted to Herbarium Women University of Azad Jammu and Kashmir, Bagh (HWAJK) for future reference. Morphologically, all specimens classified into five species i.e. *M. arvensis* (13 specimens), *M.royleana* (12 specimens), *M.piperita* (12 specimens), *M.spicata* (3 specimens) and *M. longifolia* (1 specimen) as in Fig 1. A-E.

**Fig 1.**
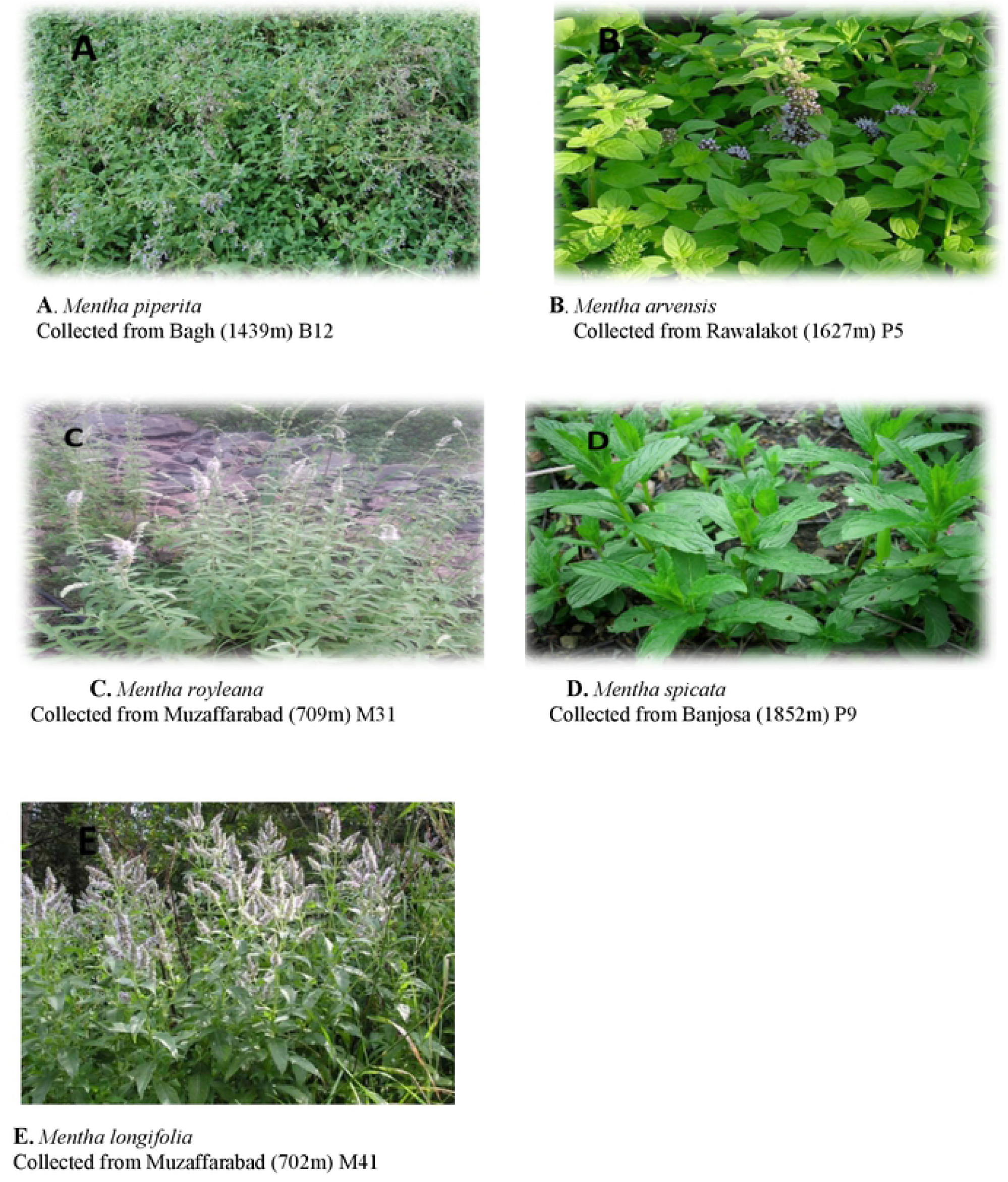
Five Mentha species identified from Azad Jammu and Kashmir A-E.

### Numerical Traits analysis

The data/plant were recorded in three replications. Observations documented for leaf area (LA), stem length (SL) and petiole length (PL) under quantitative category. Whereas, qualitative traits include leaf margin (LM), leaf apex (LAP), leaf base (LB), leaf color (LC), leaf arrangement (LAR), leaf venation (LV), leaf odor (LO), stem color (SC) and flower color (FC) stem hairiness (SH) were selected for morphological characterization. The morphological data is analyzed using SPSS software (version 23).

### Genomic DNA Extraction and DNA amplification

Genomic DNA was extracted from dried leaves using CTAB method [6]. A total of fourteen Scot and three SSR primers were used as shown in table 2. Amplification mixture for PCR reaction contained 10.5 µl of water nuclease-free, 12.5 µl of Master Mix (MBI, Fermentas), and 1µl of primer in a total of 25µl reaction volume. PCR was performed in thermal cycler with an initial denaturation temperature of 94°C for 4 minute followed by denaturation temperature 94°C for 1 minute, annealing temperature 52°-60°C for 1 minute and extension temperature of 72°C for 2 minutes Fig 2.

**Fig 2.**
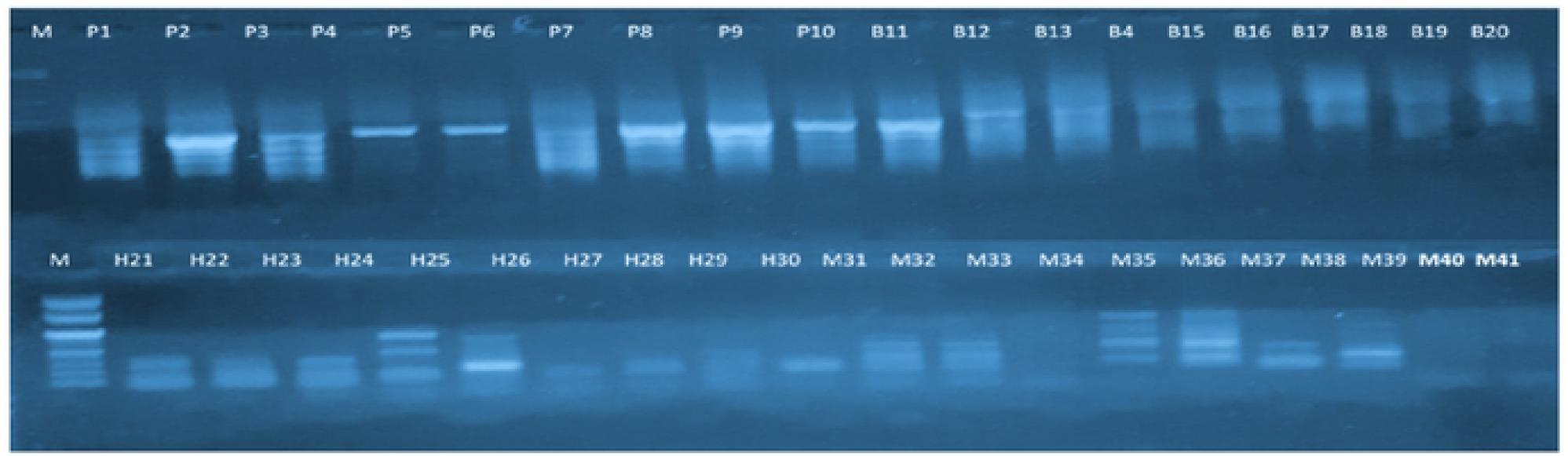
Representative gel picture of SCoT-2 primer showing polymorphic bands.

### Data Analysis

For molecular analysis data were recorded as 0 and 1 for presence or absence of bands. 0 for absence of bands and 1 for presence. Binary data were used for further data analysis, the band-based approach was used for the analysis based on [19]. The UPGMA method was used for cluster analysis and the NTSYS-pc version 2.20 was also used to determine the similarity among the *Mentha* specimens [8].

## RESULTS AND DISCUSSION

Genus *Mentha*, a taxonomically complex section, was recently studied by [18] by using RAPD fingerprinting. Genetic variations and genetic relations were evaluated among five *Mentha* taxa (*M. arvensis, M. spicata, M. royleana, M. piperita*, and *M*. longifolia). Fenwick and Ward were first to assess the genetic variations within the most commonly grown peppermint (M. piperita) and (M. gracilis) Scotch spearmint cultivars by using RAPD markers [20]. Different species of Mentha are resources for essential oils enriched in certain monoterpenes and are commonly used in food, flavor, cosmetic, and pharmaceutical industries [10]. Genus *Mentha* is complex and diversity in the taxonomy has been have been planned in past [4]. Taxonomy of Mentha is complex due to variations at morphogenetic level and presence of volatile compounds [3]. Current research study was accompanied with objectives to estimate morphogenetic variations among the *Mentha* specimens collected from different localities of Azad Jammu and Kashmir.

### Morphological study

Results of assessment between the 13 morphological tools, using statistical analyses, showed that the plant height, stem length and inflorescence variables had maximum variations among *Mentha* species, respectively. Morphological studies were carried out based on 13 morphological characters including stem length (SL), leaf area (LA), and petiole length (PL) under quantitative category. Whereas, qualitative traits include leaf margin (LM), leaf apex (LAP), leaf base (LB), leaf color (LC), leaf arrangement (LAR), leaf venation (LV), leaf odor (LO), stem color (SC) and flower color (FC) stem hairiness (SH) were selected for morphological characterization. The data was analyzed by UPGMA cluster analysis based on similarity level and using software Minitab. The 41 specimens belonging to five different species formed 4 major clusters. Group A represent M. arvensis, group B contained M.royleana and M. longifolia whereas group C contained all same species of M. piperita and group C contained M.spicata.

On the basis of morphological data *M. royleana* showed similarity with M. longifolia. Fig. 4 shows dendrogram showing morphological relationship among five species i.e., *Mentha spicata, Mentha piperita, Menthe longifolia and Mentha royleana.* On the basis of similarity coefficient, four distinct groups were observed i.e. Group A represents *M. arvensis* and Group B represents *M. royleana* respectively. It was interesting to note that each intact larger group contained samples of single species. This indicated that the status of the species is valid. Group 1 contained all samples from *Mentha arvensis* and group 2 contained all the samples of *Mentha royleana.* Dendrogram showing morphological variations among the Mentha species accordance with the previous report by Z.K. Shinwari on the basis of morphological characters [4]. According to numerical traits analysis, it is observed that numerical classification can be limited for identification of *Mentha* species having continuous polyploidy, so molecular techniques are best for proper identification of species and moreover, DNA chloroplast technology could classify upto varietal level.

**Fig 3.**
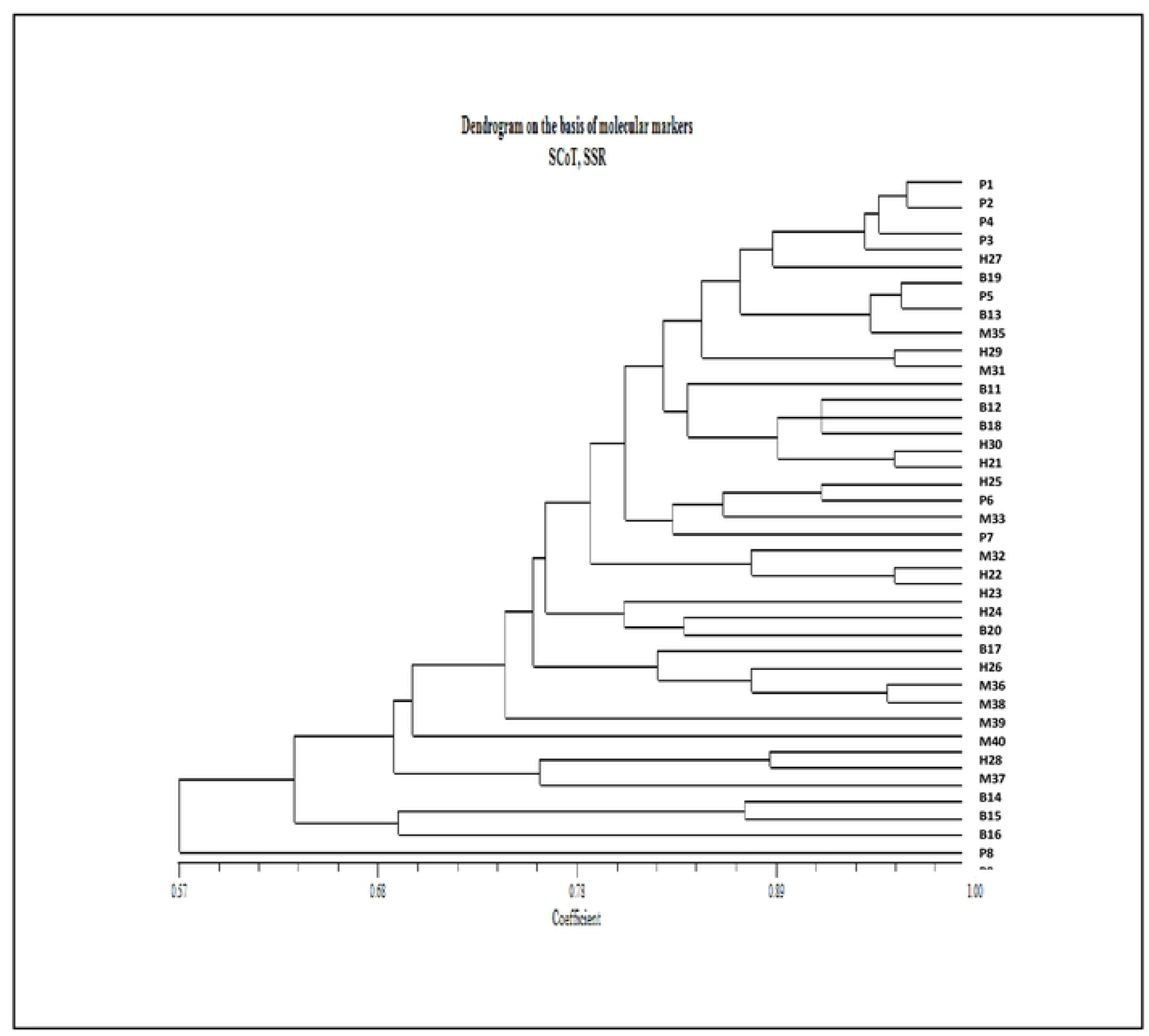
Dendrogram representing the genetic relationships among all *Mentha* genotypes using NTYSYS cluster analysis generated by fourteen molecular primers. Key: P3 *M. arvensis*, M38 *M.arvensis*, H23 *M.arvensis*, M32 *M. arvensis*, B17 *M.arvensis*, B11 *M.arvensis*, B15 *M.arvensis*, H29 *M.arvensis*, M34 *M.arvensis*, M39 *M.arvensis*, M40 *M. arvensis*, P5 *M.arvensis*, PS *M. arvensis*, H30 *M.royleana*, P2 *M. royleana*, H25 *M. royleana*, H27 *M.royleana*, H21 *M. royleana*, B16 *M.royleana*, B20 *M.royleana*, B18 *M.royleana*, H28 *M.royleana*, M3I *M.royleana*, M4I *M.longifolia*, H26 *M. royleana*, B14 *M.royleana*, B19 *M.royleana*, H22 *M.piperila*, M33 *M.piperila*, P9 *M.piperita*, Pl *M.piperita*, M37 *M.piperita*, P6 *M. piperita*, B12 *M. piperita*, P4 *M. piperita*, H24 *M.piperita*, PIO *M. piperita*, B13 *M. piperita*, M35 *M.spicata*, M36 *M.spicata*, P7 *M.spicata.*

**Fig 4.**
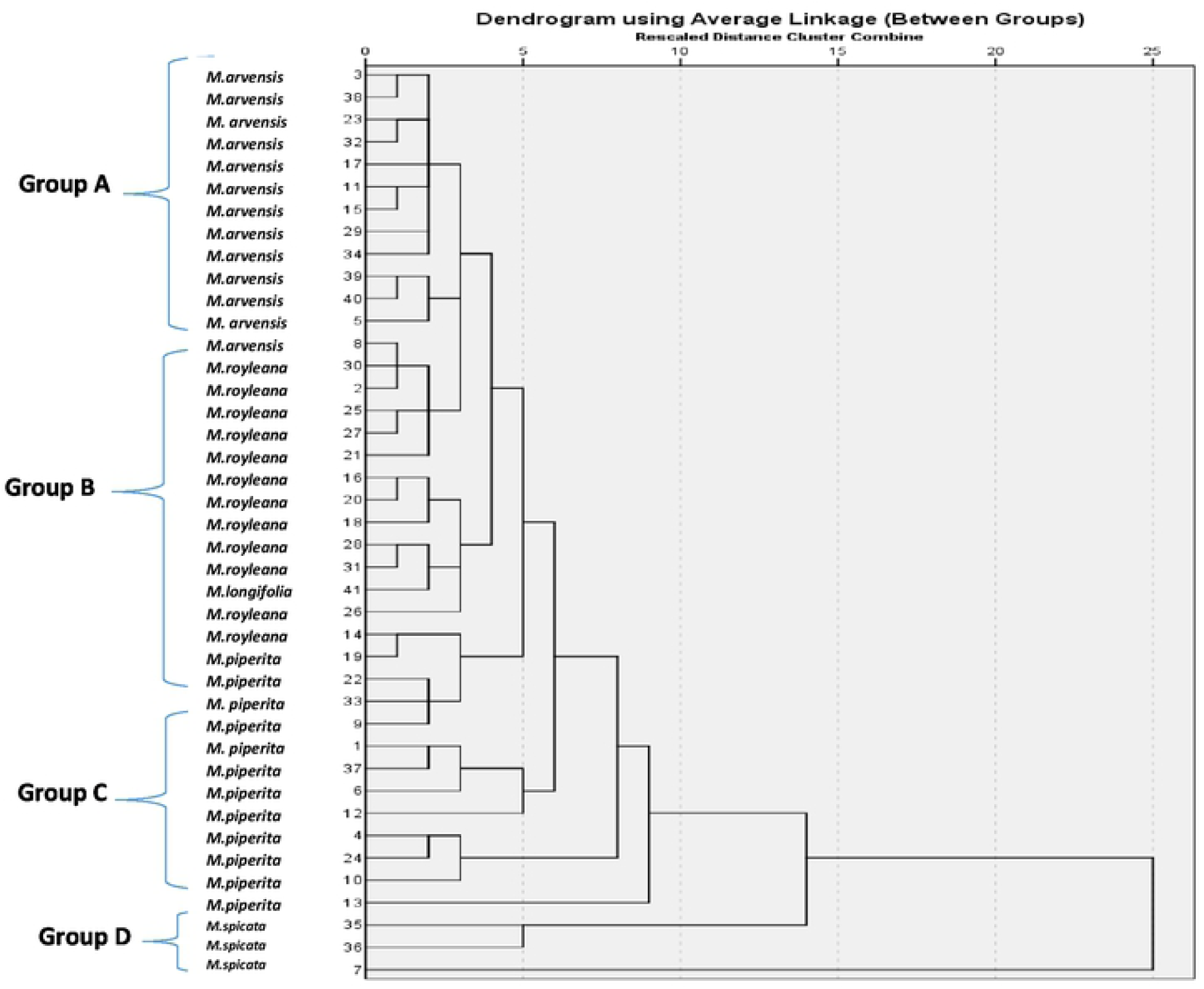
Dendrogram based on numerical traits data obtained from *Mentha* specimens, computed through SPSS program. **Key:** P3 *M. arvensis*, M38 *M.arvensis*, H23 *M.arvensis*,M32 *M.arvensis*, B17 *M.arvensis*,B11 *M.arvensis*, 815 *M.arvensis*, H29 *M.arvensis*, M34 *M.arvensis*, M39 *M.arvensis*, M40 *M.arvensis*, PS *M.arvensis*, P8 *M.arvensis*, H30 *M.royleana*, P2 *M.royleana*, H25 *M.royleana*, H27 *M.royleana*, H21 *M. royleana*, B16 *M.royleana*, B20 *M.royleana*,B18 *M.royleana*, H28 *M.royleana*, M31 *M.royleana*, M 41 *M.longifolia*, H26 *M.royleana*, B14 *M.royleana*, B19 *M.royleana*, H22 *M.piperita*, M33 *M.piperita*, P9 *M.piperita*, Pl *M.piperita*, M37 *M.piperita*, PG*M.piperita*, B12*M.piperita*, P4 *M. piperita*, H24 *M.piperita*, P10 *M.piperita*, B13 *M.piperita*, M35 *M.spicata*, M36 *M.spicata*, P7 *M.spicata.*

### Molecular study

Molecular analysis of forty one genotypes by using SCoT and SSR markers showed great genetic variability. Out of seventeen molecular markers fourteen markers produced scorable bands in forty one *Mentha* specimens. The present research is in accordance with the previous report used five RAPD primers to check diversity in thirty *Mentha* specimens [4]. Mean genetic distance estimated among 41 *Mentha* genotypes based upon the amplification pattern of SCoT and SSR primers as shown in Table 2. The mean genetic distance (GD) was calculated by computer package NTSYS version 2.02 using Dice method. Genetic distance calculated in the range of 0.35-1. The minimum GD recorded in one combination as P4-P2 while maximum GD was calculated in combination M41-P10. Estimation of genetic distance in the range of 0.35-1 is in accordance with the earlier report recorded in the range of 0.78-0.69. The present result revealed that beside morphological diversity among *Mentha* specimens collected from study area also showed high level of molecular diversity.

Overall mean Polymorphic information content (PIC) for markers calculated as 74%. The minimum PIC value by SCoT24 (57%) and the highest PIC value was recorded by SCoT 2 (80%). Major allelic frequency (MAF) was recorded in the range of 0.26 (SCoT 11) and 0.56 (tsueNH044), while mean MAF of all molecular markers was 0.36 Table 3. Polymorphic information content calculated for primers in the range from 0.57 to 0.80 per primer. PIC value confirmed that SCoT primers is efficient to evaluate the genetic diversity of medicinal plants. The present study shows SCoT primers is good alternative to check variations at molecular level in medicinal plants.

A dendrogram based on similarity coefficients generated using the weighted pair group method of arithmetic means (UPGMA) by NTSYS software presented in Fig 3. All the specimens showed maximum diversity. Specimen M41 (*M. longifolia*) recorded maximum genetic diversity and Specimen P8 (*M.arvensis*) and P9 (*M.piperita*) grouped in the same cluster belong to same species *M. arvensis* while the minimum genetic diversity estimated in P1 (*Mentha piperita*) and P2 (*Mentha royleana*). Results indicated that genus *Mentha* showed high similarity and low genetic change among different *Mentha* species.

## Conclusion

Results showed that genus *Mentha* species have great resemblance and genetic diversification among different selected species. This study presents sufficient and reliable information for successful application of SCoT and SSR technique in molecular and genetic characterization of the Mentha genotypes, as well as for variety identification. It is also evident from the PCR based assays, that SCoT and SSR-markers can be used efficiently to estimate the genetic variations among Mentha species. Maximum GD was confirmed in two combination as P4-P2, and M35-B12. So theses genotypes can further used for breeding programs. More morphological traits are required refining the data more up to varietal level. Mitochondrial and protoplast DNA technologies could be used for proper classification at genetic level.

## Acknowledgment

The author is gratefully acknowledge the kind assistance provided by Department of Genetics Hazara University Mansehra KPK, Pakistan and Department of botany Women university of Bagh AJ&K, Pakistan.

